# HTS-Oracle X: AI-Guided Prospective Discovery of Small Molecule Immune Checkpoint Binders

**DOI:** 10.64898/2026.06.17.732853

**Authors:** Somaya A. Abdel-Rahman, Moustafa Gabr

**Affiliations:** Department of Radiology, Molecular Imaging Innovations Institute (MI3), Weill Cornell Medicine, New York, NY 10065, USA

**Keywords:** Machine learning, cross-attention, immune checkpoints, virtual screening, hit discovery

## Abstract

Targeting immune checkpoint protein-protein interactions (PPIs) using small molecules remains limited by the shallow, featureless binding surfaces of co-stimulatory and co-inhibitory receptors and the characteristically low hit rates of conventional high-throughput screening against these interfaces. Here we report HTS-Oracle X, a multimodal deep learning platform that integrates bidirectional cross-attention fusion of ChemBERTa SMILES embeddings with extended RDKit descriptors, trains on continuous biophysical binding signals rather than binary labels, and employs Monte Carlo Dropout uncertainty quantification for uncertainty-adjusted compound selection. Trained on 45,760 Dianthus TRIC-screened compounds per target under scaffold-aware cross-validation, HTS-Oracle X was applied prospectively to a 100,160-compound Enamine library against CD28, TIM-3, and VISTA. From 150 model-selected compounds, 45 dose-response confirmed binders were identified (30.0% overall hit rate), yielding enrichment factors of 234-408× over experimentally established random prospective baselines and 16 sub-micromolar hits. The top hits, **HX-CD28-1** (K_D_ = 233 nM), **HX-TIM3-1** (K_D_ = 249 nM), and **HX-VISTA-1** (K_D_ = 345 nM), demonstrated on-target functional activity in immune cell and tumor co-culture assays. HTS-Oracle X represents a scalable AI-guided framework for small molecule discovery against non-enzymatic immune checkpoint targets.

**Table of Contents artwork:** 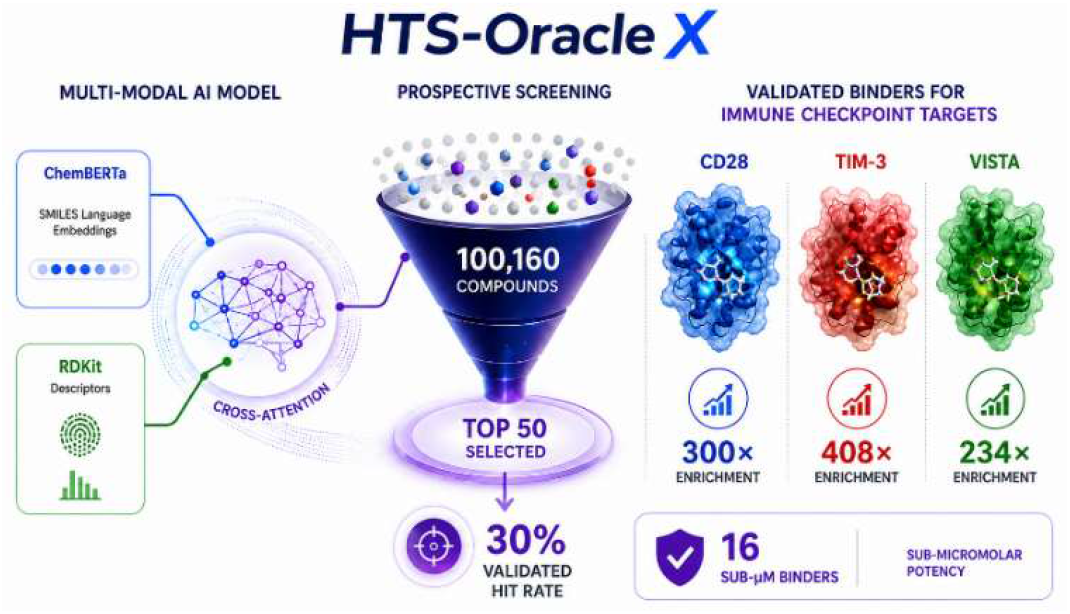

---

Immune checkpoint proteins have emerged as validated therapeutic targets across oncology, with the approval of relatlimab/nivolumab for LAG-3/PD-1 co-blockade and ongoing clinical evaluation of anti-TIGIT antibodies establishing the translational relevance of this target class beyond PD-1.^1–3^ Beyond PD-1/PD-L1, a growing number of co-stimulatory and co-inhibitory receptors have been identified as clinically actionable targets whose combined blockade or activation can overcome therapeutic resistance and broaden the scope of cancer immunotherapy. Among the next generation of immune checkpoint targets, CD28, TIM-3, and VISTA have attracted particular interest owing to their non-redundant roles in T cell activation, exhaustion, and myeloid-mediated immune suppression, respectively, yet remain underexplored by small molecule approaches relative to their clinical potential.^3,4^ Nevertheless, the broad, shallow protein-protein interaction (PPI) surfaces that mediate immune checkpoint signaling impose severe constraints on small molecule affinity and selectivity, and conventional high-throughput screening (HTS) against these interfaces routinely yields low hit rates, imposing substantial resource burdens with limited chemical output.^5–9^

Our group has developed an extensive immune checkpoint small molecule discovery program spanning LAG-3,^10–12^ ICOS,^13,14^ TIM-3,^15^ VISTA,^16^ and CD28,^17^ using pharmacophore-based virtual screening, affinity selection mass spectrometry, and antibody pharmacophore-derived hit identification workflows.^18^ These efforts provided the chemical and biological foundation for HTS-Oracle, a multimodal deep learning platform combining ChemBERTa SMILES embeddings with RDKit descriptors that achieved up to 176fold enrichment for TREM2 and CHI3L1, and 8.4-fold enrichment for CD28.^19^ However, the original platform employed simple feature concatenation, binary hit classification, and random cross-validation splits, limiting both predictive accuracy and the interpretability of prospective performance estimates.^19^

Machine learning frameworks combining molecular language models with cheminformatics features have demonstrated strong bioactivity prediction performance across diverse drug discovery applications.^20–26^ A critical but frequently overlooked requirement for platforms claiming prospective utility is scaffold-aware cross-validation, which enforces structural separation between training and test sets and prevents the performance inflation that arises from random splits when structurally similar compounds appear in both partitions.^19^ Beyond classification, training on continuous biophysical binding signals rather than binary labels preserves quantitative information that can improve the rank-ordering of active compounds, particularly in challenging PPI contexts where hit rates are low and graded binding information is diagnostic.

Here we introduce HTS-Oracle X, which advances our HTS-Oracle platform^19^ through three architectural innovations: bidirectional cross-attention fusion between the ChemBERTa and RDKit branches, regression on continuous Dianthus normalized fluorescence signal (ΔF_norm_) binding signals, and Monte Carlo Dropout uncertainty estimation enabling uncertainty-adjusted compound selection. Trained on 45,760 biophysically screened compounds per target under scaffold-aware cross-validation and prospectively applied to a 100,160-compound Enamine library, HTS-Oracle X achieves enrichment factors of 234-408× over experimentally established random prospective baselines, with 26-34% validated hit rates and sub-micromolar binders for all three targets, including a 36-fold improvement in enrichment for CD28 over the original platform.

Training datasets were established specifically for this study by our Dianthus TRIC biophysical screening of 45,760 Enamine compounds per target against recombinant CD28, TIM-3, and VISTA extracellular domains (Figures S1-S3), yielding 1,734 CD28 binders (3.79%), 2,147 TIM-3 binders (4.69%), and 1,382 VISTA binders (3.02%), with continuous ΔF_norm_ binding signals retained as regression targets rather than binary hit/non-hit labels (Figure 1A). HTS-Oracle X generates molecular representations through two parallel branches: a ChemBERTa branch producing 768-dimensional CLS token embeddings from SMILES, and an extended RDKit branch encoding Morgan fingerprints (2,048-bit), MACCS keys (167-bit), topological torsion fingerprints (1,024-bit), and 25 physicochemical descriptors (3,264 features total; Figure 1C). A bidirectional cross-attention module enables each branch to contextually attend to the other before the regression head, which predicts continuous ΔF_norm_ values directly from molecular structure (Figure 1C). Monte Carlo Dropout (10 inference passes) provides per-compound uncertainty estimates, enabling selection by Score = Predicted ΔF_norm_ − 0.5 × Uncertainty (Figure 1C). Fifteen sub-models (3 feature selection methods × 5 scaffold-aware folds) are ensemble-averaged for final predictions, with ChemBERTa embeddings pre-computed once across all compounds for efficiency (Figure 1B). The trained ensemble was applied prospectively to a 100,160-compound Enamine library, with the top 50 compounds per target selected by uncertainty-adjusted Selection Score for experimental validation (Figure 1D).

**Figure 1.**
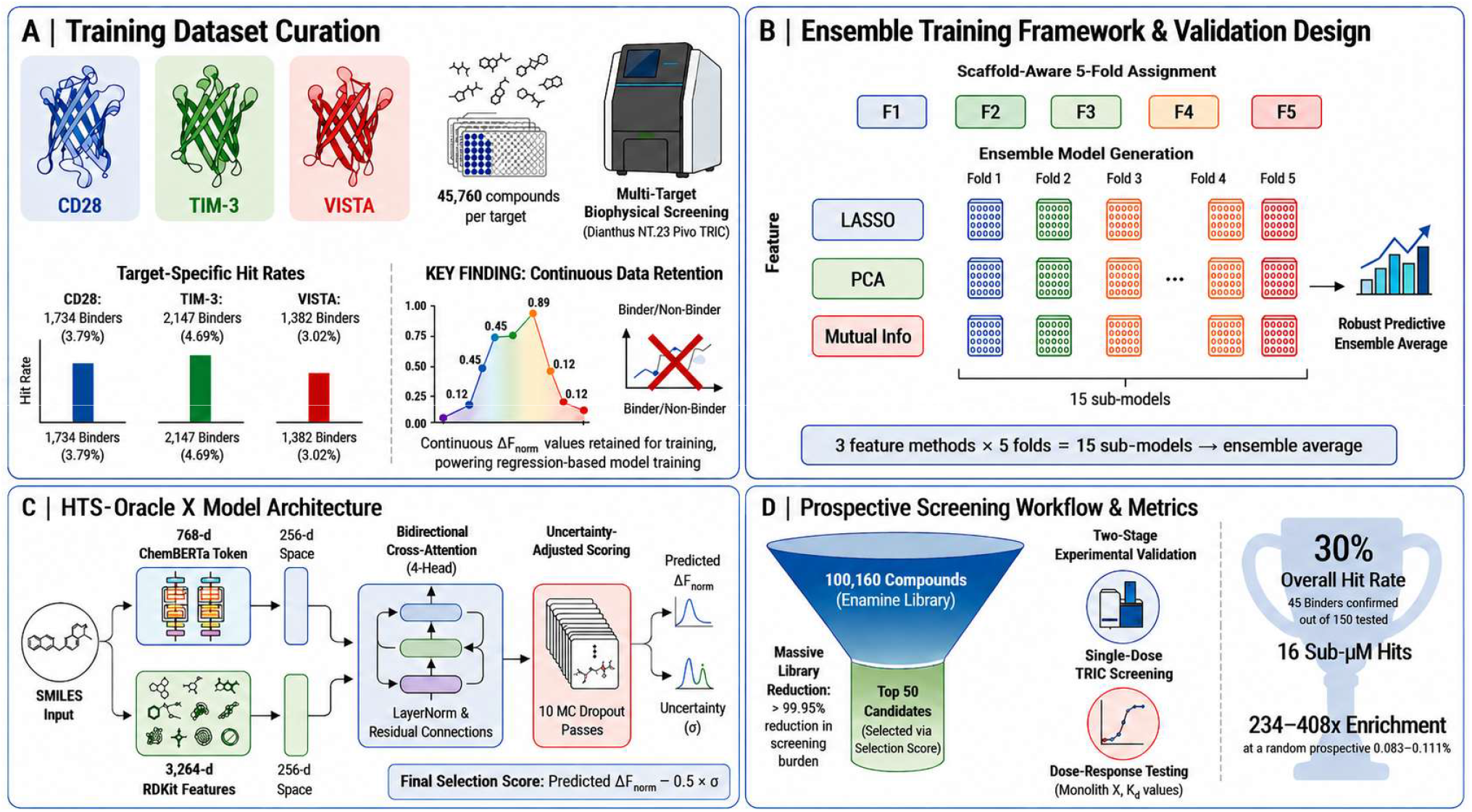
Overview of the HTS-Oracle X platform for AI-guided biophysical high-throughput screening. **(A)** Training dataset curation: 45,760 compounds per target screened by Dianthus NT.23 Pico TRIC against CD28, TIM-3, and VISTA, yielding 1,734 (3.79%), 2,147 (4.69%), and 1,382 (3.02%) confirmed binders, respectively; continuous ΔF_norm_ values are retained as regression targets rather than binary hit/non-hit labels. **(B)** Ensemble training framework: scaffold-aware 5-fold assignment distributes compounds across five folds; three feature selection methods (LASSO, PCA, Mutual Information) are applied independently across all five folds, generating 15 sub-models averaged into a robust predictive ensemble. **(C)** HTS-Oracle X architecture: ChemBERTa transformer embeddings (768-d CLS token) and extended RDKit features (3,264-d) are independently projected to 256-d representations and fused via bidirectional cross-attention (4-head, LayerNorm, residual connections); a regression head predicts continuous ΔF_norm_ binding signals, and Monte Carlo Dropout (10 passes) generates per-compound uncertainty estimates (σ) for uncertainty-adjusted Selection Score ranking (ΔF_norm_ − 0.5 × σ). **(D)** Prospective screening workflow: the 100,160-compound Enamine library is scored per target and filtered to the top 50 compounds by Selection Score (>99.95% screening burden reduction), followed by two-stage experimental validation (single-dose Dianthus TRIC then Monolith X dose-response), yielding 234-408× enrichment over random prospective baselines (0.083–0.111%), 30.0% overall hit rate (45/150 confirmed), and 16 sub-μM binders.

All performance reporting employed scaffold-aware fivefold cross-validation. HTS-Oracle X achieved ROC-AUC values of 0.914 ± 0.038 (CD28), 0.931 ± 0.032 (TIM-3), and 0.886 ± 0.049 (VISTA), with Spearman R of 0.582 ± 0.071, 0.621 ± 0.064, and 0.512 ± 0.083, respectively (Table 1; Figure 2). All targets exceeded the ROC-AUC threshold of 0.85 considered indicative of useful virtual screening performance.

**Table 1.** Computational performance of HTS-Oracle X under five-fold cross-validation.

| Target | Training Compounds | Binders (%) | ROC-AUC (mean ± SD) | Spearman R (mean ± SD) | Avg. Precision (mean ± SD) |
| --- | --- | --- | --- | --- | --- |
| CD28 | 45,760 | 1,734 (3.79%) | $0.914 \pm 0.038$ | $0.582 \pm 0.071$ | $0.742 \pm 0.052$ |
| TIM-3 | 45,760 | 2,147 (4.69%) | $0.931 \pm 0.032$ | $0.621 \pm 0.064$ | $0.778 \pm 0.047$ |
| VISTA | 45,760 | 1,382 (3.02%) | $0.886 \pm 0.049$ | $0.512 \pm 0.083$ | $0.681 \pm 0.061$ |

**Figure 2.**
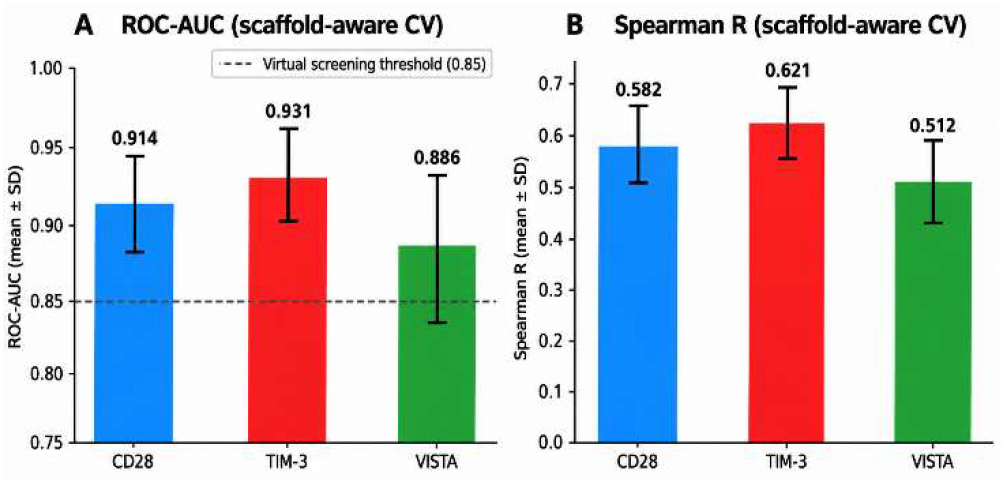
Computational performance of HTS-Oracle X under five-fold cross-validation. **(A)** ROC-AUC and **(B)** Spearman R across CD28, TIM-3, and VISTA (mean ± SD, 5 scaffold-aware folds × 3 feature methods = 15 sub-models per target). Dashed line in **(A)** indicates the ROC-AUC = 0.85 virtual screening utility threshold.

HTS-Oracle X was applied prospectively to 100,160 compounds from the Enamine Hit Locator Library, generating uncertainty-adjusted ΔF_norm_ binding score predictions independently for each immune checkpoint target. The top 50 compounds per target were selected by Selection Score (Predicted ΔF_norm_ − 0.5 × σ) and purchased for experimental validation (Figure S4), representing a 99.95% reduction in screening burden. To establish rigorous prospective enrichment baselines, 900-1,200 compounds were randomly selected from the same library and screened under identical Dianthus TRIC conditions, yielding validated hit rates of 0.100% (CD28), 0.083% (TIM-3), and 0.111% (VISTA), consistent with the characteristically low small molecule hit rates against PPI-driven immune checkpoint interfaces.

Validation of the 50 model-selected compounds per target yielded primary TRIC hit rates of 44% (CD28), 50% (TIM-3), and 40% (VISTA). Following dose-response confirmation, 15, 17, and 13 compounds were validated as confirmed binders (30%, 34%, and 26% hit rates), corresponding to enrichment factors of 300×, 408×, and 234× over the random prospective baselines (Table S1; Figure 3). Attrition of 31.8-35.0% from primary TRIC to confirmed binding reflects rigorous false positive exclusion. Across all three targets, 45 of 150 model-selected compounds were confirmed binders (30.0% overall hit rate).

**Figure 3.**
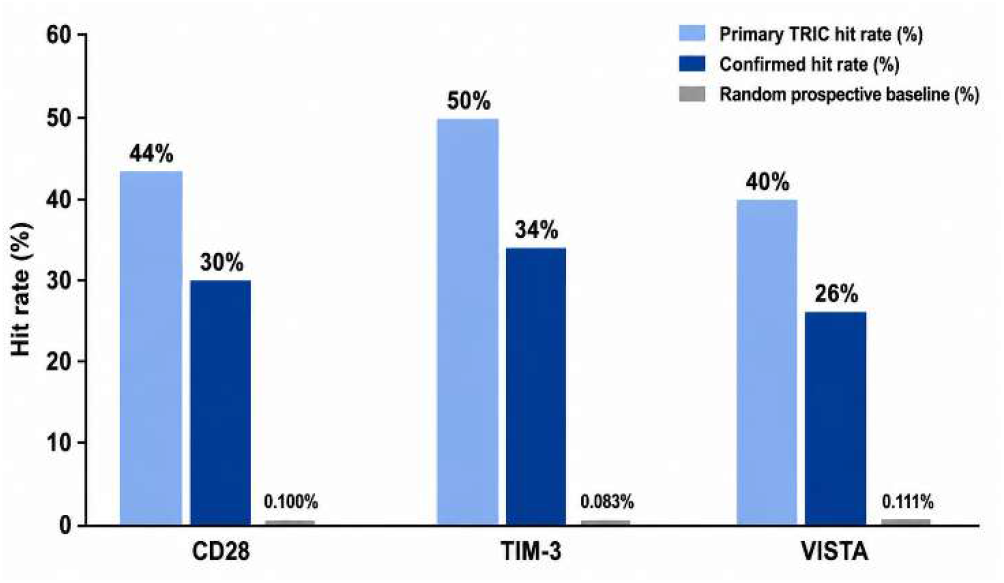
Prospective validation hit rates for HTS-Oracle X. Primary TRIC hit rates (light blue), dose-response confirmed hit rates (dark blue), and experimentally established random prospective baseline hit rates (grey) for CD28, TIM-3, and VISTA. Confirmed hit rates of 26-34% correspond to enrichment factors of 234-408× over the random prospective baseline

Among 45 confirmed hits (Table S2), 16 demonstrated sub-micromolar equilibrium dissociation constant (K_D_) values: 7 for CD28, 7 for TIM-3, and 2 for VISTA (Table 2; Figure 4). For CD28, top hits **HX-CD28-1** (K_D_ = 0.233 µM), **HXCD28-2** (K_D_ = 0.262 µM), and **HX-CD28-3** (K_D_ = 0.443 µM) extend the chemical diversity of CD28 small molecule binders reported by our group.^17^ For TIM-3, **HX-TIM3-1** (K_D_ = 0.249 µM), **HX-TIM3-2** (K_D_ = 0.257 µM), and **HX-TIM3-3** (K_D_ = 0.289 µM) complement TIM-3 binders identified through pharmacophore-based approaches.^15^ For VISTA, **HX-VISTA-1** (K_D_ = 0.345 µM) represents a remarkable addition to the limited small molecule chemical matter available for this target.^16,18^

**Table 2.** Top confirmed binders per target from HTS-Oracle X prospective screening.

| Target | Compound ID | $K_D$ ($\mu\text{M}$) | Pred. $\Delta F_{\text{norm}}$ (%) | Selection Score |
| --- | --- | --- | --- | --- |
| CD28 | **HX-CD28-1** | 0.233 | 14.43 | 14.20 |
| CD28 | **HX-CD28-2** | 0.262 | 20.37 | 19.96 |
| CD28 | **HX-CD28-3** | 0.443 | 17.94 | 17.64 |
| TIM-3 | **HX-TIM3-1** | 0.249 | 17.13 | 16.72 |
| TIM-3 | **HX-TIM3-2** | 0.257 | 20.15 | 19.76 |
| TIM-3 | **HX-TIM3-3** | 0.289 | 16.04 | 15.52 |
| VISTA | **HX-VISTA-1** | 0.345 | 13.35 | 12.77 |
| VISTA | **HX-VISTA-2** | 0.931 | 16.46 | 15.71 |
| VISTA | **HX-VISTA-3** | 1.004 | 14.80 | 14.28 |

**Figure 4.**
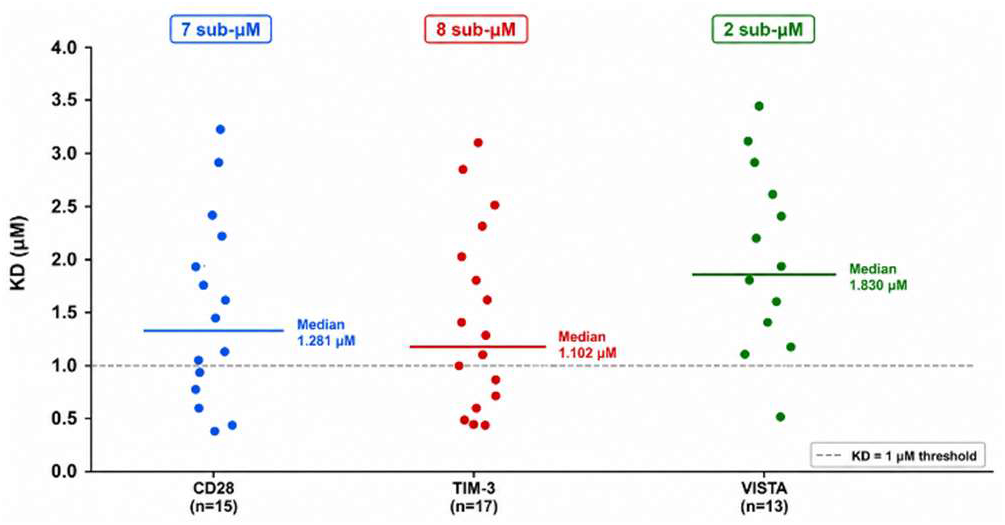
Binding affinity (K_D_) distribution of confirmed hits per immune checkpoint target. Individual data points represent confirmed binders; horizontal lines indicate median K_D_ values. Numbers in boxes indicate sub-micromolar hits per target. K_D_ values determined by Monolith X spectral shift dose-response.

Enrichment factors of 300× (CD28), 408× (TIM-3), and 234× (VISTA) were calculated against experimentally established random prospective baselines from the same 100,160-compound library (Figure 5A). This approach, in which the enrichment denominator is derived from experimental random screening of the same prospective library, provides the most rigorous and scientifically defensible enrichment estimate, directly comparable across platforms and target classes. For CD28, the only target shared with the original HTS-Oracle platform,^19^ HTS-Oracle X achieves 300× enrichment vs 8.4× for the original platform, a 36-fold improvement, alongside a step change in screening burden reduction from 70% to 99.95% and a substantially larger prospective screening library (100,160 vs 1,152 compounds; Figure 5B). The most potent CD28 binder (**HX-CD28-1**, K_D_ = 0.233 µM) demonstrates sub-micromolar target engagement, extending the affinity frontier for small molecule CD28 modulators.

**Figure 5.**
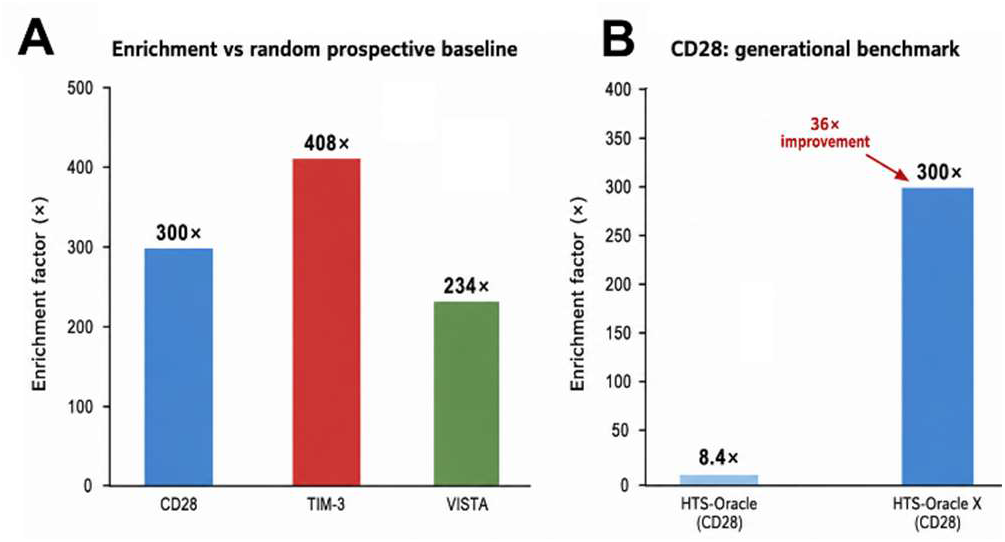
Enrichment factors for HTS-Oracle X. **(A)** Enrichment factors for CD28 (300×), TIM-3 (408×), and VISTA (234×) calculated against experimentally established random prospective baselines from the same 100,160-compound library. **(B)** Generational comparison for CD28: HTS-Oracle (8.4×) vs HTS-Oracle X (300×), representing a 36-fold improvement.

To characterize the structural and functional properties of the most potent confirmed binders, the top hit compound per target was advanced to comprehensive biophysical and cellular evaluation (Figure 6). The chemical structures of **HX-CD28-1, HX-TIM3-1**, and **HX-VISTA-1** reveal structurally distinct small molecule scaffolds across the three targets, reflecting the chemical diversity captured by HTS-Oracle X across the Enamine library (Figures 6A–C). Doseresponse binding isotherms obtained by Monolith X spectral shift confirmed sub-micromolar affinities for all three compounds: **HX-CD28-1** bound CD28 with a K_D_ of 233 ± 24.7 nM (Figure 6D), **HX-TIM3-1** bound TIM-3 with a K_D_ of 249 ± 16.3 nM (Figure 6E), and **HX-VISTA-1** bound VISTA with a K_D_ of 345 ± 46.1 nM (Figure 6F), collectively establishing direct and target-selective engagement for all three scaffolds. These affinities are among the highest reported for small molecule binders of CD28, TIM-3, and VISTA extracellular domains and represent a meaningful advance in the available chemical matter for this target class.

**Figure 6.**
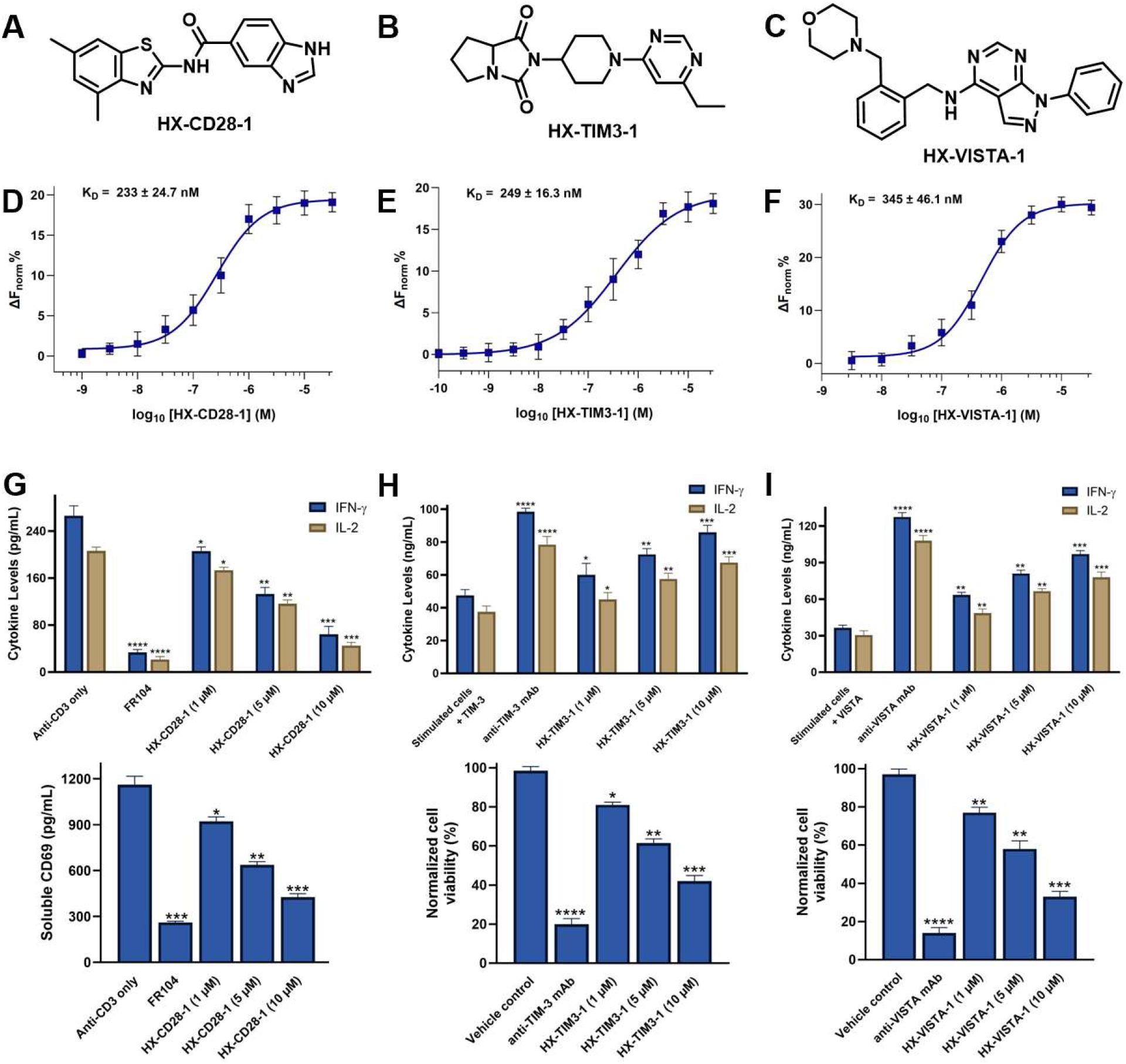
Chemical structures, biophysical binding, and functional characterization of top hit compounds identified by HTS-Oracle X for CD28, TIM-3, and VISTA. **(A–C)** Chemical structures of **HX-CD28-1 (A), HX-TIM3-1 (B)**, and **HX-VISTA-1 (C). (D-F)** Monolith X binding isotherms for **HX-CD28-1** binding to CD28 (**D**, K_D_ = 233 ± 24.7 nM), **HX-TIM3-1** binding to TIM-3 (**E**, K_D_ = 249 ± 16.3 nM), and **HX-VISTA-1** binding to VISTA (**F**, K_D_ = 345 ± 46.1 nM). Data are presented as mean ± SD (n=5). **(G)** Functional evaluation of **HX-CD28-1** in a tumor-PBMC co-culture assay. IFN-γ and IL-2 secretion (top) and soluble CD69 levels (bottom) were measured after 48-hour co-culture of A549 tumor spheroids with human PBMCs (E:T ratio 5:1) in the presence of anti-CD3 (0.3 µg/mL), FR104 (10 µg/mL, positive control), or **HX-CD28-1** (1, 5, 10 µM). **(H)** Functional evaluation of **HX-TIM3-1**. IFN-γ and IL-2 cytokine levels were measured in PBMCs cultured with recombinant TIM-3 in the presence of anti-TIM-3 mAb (positive control) or HX-TIM3-1 (1, 5, 10 µM) (top). Normalized cell viability of THP-1 AML cells, which endogenously express TIM-3, was assessed following compound treatment (bottom). **(I)** Functional evaluation of **HX-VISTA-1**. IFN-γ and IL-2 cytokine levels were measured in PBMCs cultured with recombinant VISTA in the presence of anti-VISTA mAb (positive control) or **HX-VISTA-1** (1, 5, 10 µM) (top). Normalized cell viability of SKOV3 ovarian cancer cells, which endogenously express high levels of VISTA, was assessed following compound treatment (bottom). For all bar graphs, data represent mean ± SEM of *n* = 6 independent wells. Statistical comparisons were made to the respective vehicle or stimulated-only control using one-way ANOVA with Dunnett’s posthoc test. ns, not significant; \**p* < 0.05; \*\**p* < 0.01; \*\*\**p* < 0.001; \*\*\*\**p* < 0.0001.

Functional evaluation of **HX-CD28-1** was performed in a tumor–PBMC co-culture assay using A549 tumor spheroids (E:T ratio 5:1) with anti-CD3 stimulation (Figure 6G). Treatment with **HX-CD28-1** produced dose-dependent suppression of IFN-γ and IL-2 secretion and a parallel reduction in soluble CD69 levels, a surface marker of early T cell activation, at 1, 5, and 10 μM (Figure 6G, top and bottom panels). The magnitude of suppression at 5-10 μM approached that of FR104 (10 μg/mL), a clinical-stage CD28-selective biologic antagonist used as a positive control, demonstrating that **HX-CD28-1** engages CD28 co-stimulatory signaling in a physiologically relevant cellular context. For TIM-3, **HXTIM3-1** was evaluated across two orthogonal functional readouts (Figure 6H). In a human PBMC/recombinant TIM-3 cytokine assay, **HX-TIM3-1** restored IFN-γ and IL-2 production in a dose-dependent manner relative to the TIM-3-suppressed baseline, consistent with relief of TIM-3-mediated immune suppression (Figure 6H, top panel). Complementarily, **HX-TIM3-1** reduced the viability of THP-1 acute myeloid leukemia (AML) cells, which endogenously express TIM-3, in a concentration-dependent fashion at 1-10 μM, with efficacy comparable to the anti-TIM-3 monoclonal antibody positive control at higher concentrations (Figure 6H, bottom panel). TIM-3 expression on AML blasts has been established as a driver of immune evasion and disease progression, and the dual cytokine and viability activity of **HX-TIM3-1** provides orthogonal evidence of ontarget functional engagement. An analogous two-readout functional profile was obtained for **HX-VISTA-1** (Figure 6I): dose-dependent restoration of IFN-γ and IL-2 in PBMCs cultured with recombinant VISTA (Figure 6I, top panel) and concentration-dependent reduction in viability of SKOV3 ovarian cancer cells, which endogenously express high levels of VISTA, with an efficacy profile comparable to the anti-VISTA monoclonal antibody control (Figure 6I, bottom panel).

Taken together, the structural diversity of the three hit scaffolds, their sub-micromolar biophysical affinities, and their on-target functional activity across mechanistically distinct immune checkpoint contexts collectively establish Figure 6 as the critical proof-of-concept validation that HTS-Oracle X identifies not merely biophysical binders but functionally active small molecule immune checkpoint modulators, a distinction of central importance for the translational relevance of AI-guided hit discovery platforms in immunooncology.

In summary, HTS-Oracle X establishes a new performance benchmark for AI-guided small molecule discovery against immune checkpoint PPIs. Three architectural advances over the original HTS-Oracle platform, bidirectional crossattention fusion, continuous ΔF_norm_ regression, and Monte Carlo Dropout uncertainty quantification, collectively drive a 36-fold improvement in CD28 enrichment and deliver 234-408× prospective enrichment factors across all three targets, with 45 dose-response confirmed binders from 150 model-selected compounds (30.0% overall hit rate) and 16 sub-micromolar hits. Critically, the top hit per target, **HX-CD28-1** (K_D_ = 233 nM), **HX-TIM3-1** (K_D_ = 249 nM), and **HX-VISTA-1** (K_D_ = 345 nM), not only confirm direct sub-micromolar target engagement by Monolith X but also demonstrate on-target functional activity across mechanistically distinct immune cell and tumor co-culture assays, establishing that HTS-Oracle X delivers functionally validated chemical matter rather than biophysical binders alone. Importantly, the platform is generalizable to any immune checkpoint for which biophysical screening data are available. While the present study focused on biophysical enrichment and initial functional validation, future work will include medicinal chemistry optimization, selectivity profiling, and in vivo pharmacologic characterization

## Supporting Information

The Supporting Information is available free of charge on the ACS Publications website.

Experimental procedures, prospective validation results for HTS-Oracle X, complete list of 45 dose-response confirmed binders identified by HTS-Oracle X, primary Dianthus TRIC biophysical screening against CD28, TIM-3, and VISTA, and selection score distributions for the three targets (PDF)

## Author information

### Author Contributions

The manuscript was written through contributions of all authors. All authors have given approval to the final version of the manuscript.

## Abbreviations

AI: artificial intelligence;
AML: acute myeloid leukemia;
AUC: area under the curve;
CD28: cluster of differentiation 28;
CLS: classification token;
ΔF~norm~: normalized fluorescence change;
E:T: effector-to-target;
HTS: high-throughput screening;
IFN-γ: interferon gamma;
IL-2: interleukin-2;
ICOS: inducible T cell costimulator;
K~D~: equilibrium dissociation constant;
LAG-3: lymphocyte activation gene 3;
LASSO: least absolute shrinkage and selection operator;
mAb: monoclonal antibody;
MCD: Monte Carlo Dropout;
MI: mutual information;
PCA: principal component analysis;
PBMC: peripheral blood mononuclear cell;
PD-1: programmed cell death protein 1;
PD-L1: programmed death-ligand 1;
PPI: protein–protein interaction;
ROC: receiver operating characteristic;
SD: standard deviation;
SEM: standard error of the mean;
SMILES: simplified molecular-input line-entry system;
TIM-3: T cell immunoglobulin and mucin domain-containing protein 3;
TIGIT: T cell immunoreceptor with Ig and ITIM domains;
TRIC: temperature-related intensity change;
VISTA: V-domain Ig suppressor of T cell activation

## NOTES

The authors declare no competing financial interests. The code is available on https://github.com/gabr2003/HTS-Oracle-X

