## Supporting Information for "HTS-Oracle X: AI-Guided Prospective Discovery of Small Molecule Immune Checkpoint Binders"

*Electronic Supplementary Information for*

AUTHOR ADDRESS

| **Contents** |  |
| --- | --- |
| Experimental  Prospective validation results for HTS-Oracle X | S3  S6 |
| Complete list of 45 dose-response confirmed binders identified by HTS-Oracle X prospective screening | S7 |
| Dianthus TRIC single-dose screening of the 45,760-compound Enamine training library against CD28 | S10 |
| Dianthus TRIC single-dose screening of the 45,760-compound Enamine training library against TIM-3 | S11 |
| Dianthus TRIC single-dose screening of the 45,760-compound Enamine training library against VISTA | S11 |
| Selection Score distributions across the 100,160-compound Enamine library for CD28, TIM-3, and VISTA | S12 |

**Experimental:**

**Compound Procurement**

All compounds evaluated in this study were commercially available compounds purchased from Enamine (Monmouth Junction, NJ, USA). Enamine IDs for all validated hits from this study are provided in Table S2.

### **Biophysical Training Dataset**

Dianthus NT.23 Pico TRIC single-dose screening (NanoTemper Technologies) of 45,760 Enamine compounds was performed against recombinant CD28, TIM-3, and VISTA extracellular domains labelled with RED-tris-NTA 2nd Generation dye (10 nM). Confirmed binders exceeded mean + 3 SD of vehicle controls. Continuous ΔF_norm_ values were retained for all compounds as regression targets.

Recombinant human His-tagged CD28, TIM-3, or VISTA protein wasfluorescently labeled with RED-tris-NTA 2nd Generation dye (NanoTemper, Cat. #MO-L018) according to the manufacturer’s protocol. Labeled protein (final concentration: 20 nM) was incubated with test compounds at a fixed concentration (10 µM) in assay buffer (PBS pH 7.4 supplemented with 0.05% Tween-20 and 1% (v/v) DMSO). Fluorescence was measured in Dianthus 384-well plates (NanoTemper Technologies, Catalog# DI-P001A) with LED power set to 40%. Fluorescence was recorded at 670 nm and 650 nm, and normalized fluorescence (F_norm_) was calculated as the ratio of F670/F650. Binding responses were quantified as F_norm_.

**Quantitative Binding Affinity Determination Using Monolith**

Binding affinities of selected hits were quantified by microscale thermophoresis using a Monolith X instrument (NanoTemper Technologies). His-tagged protein (CD28, TIM-3, or VISTA) was labeled with RED-tris-NTA 2nd Generation dye using the Monolith His-Tag Labeling Kit (Cat. #MO-L018) following the manufacturer’s guidelines. Compound titrations were prepared as serial dilutions (PBS buffer, pH 7.4, 0.05% Tween-20, 1% (v/v) DMSO). Following a 15 min incubation at room temperature in the dark, samples were loaded into Monolith capillaries (Cat. #MO-K022) and analyzed at 25 °C using 40% LED power and medium MST power settings. Normalized fluorescence (F_norm_) values were determined as the ratio of fluorescence intensity after and before IR laser heating. Each compound was evaluated in five technical replicates. Dissociation constants (K_D_) were calculated from three independent experiments using MO.Affinity Analysis software and GraphPad Prism 10, applying standard dose-response fitting models. Data represent mean ± SD (n=5).

### **Model Architecture and Training**

RDKit features (Morgan 2048-bit, MACCS 167-bit, topological torsion 1024-bit, 25 descriptors; 3,264 total) were combined with pre-computed ChemBERTa CLS embeddings (768-d) through bidirectional cross-attention (4 heads) before a 512→256→128→1 regression head. Training used Huber loss, AdamW (lr = 2×10⁻⁴), cosine annealing, and early stopping (patience = 5, Spearman R). Fifteen sub-models (LASSO/PCA/mutual information × 5 scaffold-aware folds) were ensemble-averaged. Monte Carlo Dropout (10 passes) provided uncertainty estimates.

### **Scaffold-Aware Cross-Validation**

Murcko scaffolds were computed using RDKit. Compounds with identical scaffolds were assigned to the same fold, ensuring zero structural leakage. Performance was assessed by ROC-AUC, Spearman R, and average precision from out-of-fold predictions.

### **Prospective Screening and Experimental Validation**

The 100,160-compound Enamine library was scored per target; compounds were ranked by Selection Score = Predicted ΔF_norm_ − 0.5 × Uncertainty; the top 50 were purchased. Model-selected and random baseline compounds were screened by Dianthus TRIC. Primary hits underwent Monolith X dose-response; K_D_ values were fit by nonlinear regression in GraphPad Prism 10. Enrichment factor = confirmed hit rate (model) / confirmed hit rate (random baseline). Screening burden reduction = (1 − 50/100,160) × 100%.

### **Evaluation of T Cell Activation in Tumor–PBMC Co-culture (HX-CD28-1)**

To assess the functional activity of **HX-CD28-1** in a physiologically relevant immunological context, a 3D tumor–PBMC co-culture assay was employed. A549 human lung adenocarcinoma cells (from ATCC) were seeded in ultra-low attachment 96-well plates (1,000 cells/well) and allowed to form spheroids over 48 hours. Spheroids were subsequently treated with recombinant human interferon-γ (IFN-γ, 50 ng/mL) for 24 hours to upregulate surface CD80/CD86 expression and prime the tumor–immune interface. Freshly thawed human peripheral blood mononuclear cells (PBMCs) were added to the spheroid-containing wells at a 5:1 effector-to-target (E:T) ratio. Co-cultures were stimulated with soluble anti-CD3 antibody (0.3 μg/mL, clone OKT3) to model TCR-dependent activation. **HX-CD28-1** was added at the initiation of co-culture at concentrations of 1, 5, and 10 μM. FR104 (10 μg/mL), a clinical-stage monovalent anti-CD28 Fab’ fragment, served as a positive control for CD28 blockade. Vehicle (DMSO) and anti-CD3-only conditions were included as controls. After 48 hours, culture supernatants were harvested and analyzed for secreted IFN-γ and IL-2 levels using human-specific ELISA kits (BioLegend) according to the manufacturer’s instructions. Soluble CD69 (sCD69), shed from the surface of activated T cells upon stimulation, was quantified using a human CD69 ELISA kit (ThermoFisher). All cytokine concentrations were interpolated from four-parameter standard curves. Data represent mean ± SEM of n = 6 independent wells. Statistical comparisons to the anti-CD3 only control group were performed using one-way ANOVA followed by Dunnett’s post-hoc test.

### **Assessment of TIM-3 Functional Activity: Cytokine Release and AML Cell Viability (HX-TIM3-1)**

Human T cells were isolated from PBMCs using the Human Pan-T Cell Isolation Kit (Miltenyi Biotec) and cultured in RPMI-1640 supplemented with 10% fetal bovine serum (FBS) and 1% penicillin/streptomycin. T cells were stimulated with plate-bound anti-CD3 (1 μg/mL) and soluble anti-CD28 (1 μg/mL) in the presence of recombinant human TIM-3 protein (25 nM) to model TIM-3-mediated immunosuppression. **HX-TIM3-1** was added at concentrations of 1, 5, and 10 μM. Anti-TIM-3 monoclonal antibody (M6903, 100 nM) served as a positive control for TIM-3 blockade. After 48 hours, supernatants were harvested and IFN-γ and IL-2 levels were quantified using human-specific ELISA kits (BioLegend) according to the manufacturer’s instructions. Data represent mean ± SEM of n = 6 independent wells.

THP-1 cells (from ATCCC), a human monocytic AML cell line, were cultured in RPMI-1640 supplemented with 10% FBS, 100 units/mL penicillin, and 100 μg/mL streptomycin at 37°C in a humidified 5% CO_2_ atmosphere. THP-1 cells were co-cultured with freshly thawed PBMCs at a 1:5 target-to-effector ratio. **HX-TIM3-1** was added at concentrations of 1, 5, and 10 μM. Anti-TIM-3 monoclonal antibody (M6903, 100 nM) served as the positive control. Cell viability was assessed using the CellTiter-Glo® Luminescent Cell Viability Assay (Promega) according to the manufacturer's instructions. Briefly, after 72 hours of co-culture, CellTiter-Glo® reagent was added at a 1:1 ratio to the culture medium, and luminescence was recorded using a GloMax® plate reader. Luminescent signal is proportional to ATP content and reflects the number of metabolically active cells. Data represent mean ± SEM of n = 6 independent wells. Statistical comparisons to the vehicle control group were performed using one-way ANOVA followed by Dunnett’s post-hoc test.

### **Assessment of VISTA Functional Activity: Cytokine Release and Ovarian Cancer Cell Viability (HX-VISTA-1)**

Human T cells were isolated from PBMCs using the Human Pan-T Cell Isolation Kit (Miltenyi Biotec) and cultured in RPMI-1640 supplemented with 10% FBS and 1% penicillin/streptomycin. T cells were stimulated with plate-bound anti-CD3 (1 μg/mL) and soluble anti-CD28 (1 μg/mL) in the presence of recombinant human VISTA protein (25 nM) to model VISTA-mediated T cell suppression. **HX-VISTA-1** was added at concentrations of 1, 5, and 10 μM. Anti-VISTA monoclonal antibody (100 nM) served as the positive control for VISTA blockade. After 48 hours, supernatants were harvested and IFN-γ and IL-2 concentrations were quantified using human-specific ELISA kits (BioLegend) according to the manufacturer’s instructions. Data represent mean ± SEM of n = 6 independent wells.

SKOV3 human ovarian adenocarcinoma cells (from ATCC), which endogenously express high levels of VISTA, were cultured in McCoy’s 5A medium supplemented with 10% FBS and 1% penicillin/streptomycin at 37°C in a humidified 5% CO_2_ atmosphere. SKOV3 cells were co-cultured with freshly thawed PBMCs at a 1:5 target-to-effector ratio, following the same protocol used for the TIM-3 viability assay. **HX-VISTA-1** was added at concentrations of 1, 5, and 10 μM. Anti-VISTA monoclonal antibody (100 nM) served as the positive control. Cell viability was assessed using the CellTiter-Glo® Luminescent Cell Viability Assay (Promega) according to the manufacturer's instructions. Briefly, after 72 hours of co-culture, CellTiter-Glo® reagent was added at a 1:1 ratio to the culture medium, and luminescence was recorded using a GloMax® plate reader. Luminescent signal is proportional to ATP content and reflects the number of metabolically active cells. Normalized cell viability was expressed as percentage of the vehicle-treated control. Data represent mean ± SEM of n = 6 independent wells. Statistical comparisons to the vehicle control group were performed using one-way ANOVA followed by Dunnett’s post-hoc test.

**Table S1. Prospective validation results for HTS-Oracle X.**

| **Target** | **Tested** | **Primary hits** | **Confirmed** | **Hit rate (%)** | **Random baseline (%)** | **Enrichment (×)** | **Burden reduction (%)** |
| --- | --- | --- | --- | --- | --- | --- | --- |
| CD28 | 50 | 22 (44%) | 15 | 30.0 | 0.100 | 300× | 99.95 |
| TIM-3 | 50 | 25 (50%) | 17 | 34.0 | 0.083 | 408× | 99.95 |
| VISTA | 50 | 20 (40%) | 13 | 26.0 | 0.111 | 234× | 99.95 |
| Overall | 150 | 67 (45%) | 45 | 30.0 | — | — | 99.95 |

**Table S2.** Complete list of 45 dose-response confirmed binders identified by HTS-Oracle X prospective screening, designated HX-[Target]-[Number] and sorted by ascending K_D_ within each target. K_D_ values determined by Monolith X spectral shift dose-response. All compounds purchased from Enamine.

| **Compound** | **Enamine ID** | **Target** | **SMILES** | **KD (µM)** |
| --- | --- | --- | --- | --- |
| **CD28 Confirmed Binders (n = 15)** | | | | |
| **HX-CD28-1** | Z30268329 | CD28 | CC=1C=C(C)C=2N=C(NC(=O)C=3C=CC=4NC=NC4C3)SC2C1 | 0.233 |
| **HX-CD28-2** | Z1280348291 | CD28 | CCCC(=O)N1CCCCC1C(=O)N2CCCC(C2)S(N)(=O)=O | 0.262 |
| **HX-CD28-3** | Z466958190 | CD28 | COC(=O)C=1C=CC=C(C1)C(=O)NC2CCC(CC2)C(F)(F)F | 0.443 |
| **HX-CD28-4** | Z729726666 | CD28 | CC1=NOC(=N1)NC(=O)CC=2C=CC=C3C=CC=CC23 | 0.528 |
| **HX-CD28-5** | Z3347328650 | CD28 | C[C@H](N)C=1C=CC(=CC1)NC(=O)N2CCC(C)S(=O)(=O)CC2 | 0.535 |
| **HX-CD28-6** | Z64566750 | CD28 | O=S(=O)(NCC1=CC=CO1)C=2C=CC=C(C2)NC=3N=CN=C4SC=5CCCC5C34 | 0.654 |
| **HX-CD28-7** | Z2699389052 | CD28 | CCO[C@@]1(C)C[C@H]1NC(=O)N[C@H]2CCN(C2)S(=O)(=O)C=3C=CC=CC3 | 0.966 |
| **HX-CD28-8** | Z433877472 | CD28 | COCC(=O)N1CCN(CC2=NC(=NO2)C3=CC=C(C)C(F)=C3)CC1 | 1.28 |
| **HX-CD28-9** | Z19845168 | CD28 | CSC1=NC=2C=CC=CC2C(=O)N1CC3CCCO3 | 1.72 |
| **HX-CD28-10** | Z2608261393 | CD28 | CC1=NNC(CC(=O)N2CCOCC2C=3C(C)=NN(CC=4C=CC=CC4)C3C)=N1 | 1.83 |
| **HX-CD28-11** | Z2304036986 | CD28 | CCO[C@H]1C[C@H](C1)NC(=O)C=2C(Cl)=CC(Cl)=CC2O | 5.49 |
| **HX-CD28-12** | Z1759676666 | CD28 | CC1=CC=C(C(=O)NC[C@@H]2C[C@H](F)CN2C=3C=CN=C(N3)N(C)C)C(=O)N1 | 6.86 |
| **HX-CD28-13** | Z1685733148 | CD28 | CC=1N=C(SC1C2=NC(=NO2)C3CC(=O)N(CC4CCC4)C3)C=5C=NN(C)C5 | 8.95 |
| **HX-CD28-14** | Z3016310107 | CD28 | C=CCC(C(=O)N1CCC(C1)C=2C=NC=CN2)C=3C=CC=CC3 | 10.65 |
| **HX-CD28-15** | Z1986422089 | CD28 | CCC=1C=CC=C2C(=CNC12)C3CCN(CC3)C(=O)C=4C=NC5=CN=CC=C5C4 | 11.55 |
| **TIM-3 Confirmed Binders (n = 17)** | | | | |
| **HX-TIM3-1** | Z1612723001 | TIM-3 | CCC=1C=C(N=CN1)N2CCC(CC2)N3C(=O)[C@@H]4CCCN4C3=O | 0.249 |
| **HX-TIM3-2** | Z230375366 | TIM-3 | CCOC=1C=CC(=CC1S(=O)(=O)N(C)C)NC(=O)C2CC(=O)NC=3C=C(F)C=CC32 | 0.257 |
| **HX-TIM3-3** | Z165420654 | TIM-3 | NC(=O)C=1C=CC(=CC1)CN2C=NC=3C(Cl)=CC(Cl)=CC3C2=O | 0.289 |
| **HX-TIM3-4** | Z2348165329 | TIM-3 | CC=1C=C(N(C)N1)N2CCCC(C2)NC(=O)N3CCC(CC3)C=4C=CN=CN4 | 0.355 |
| **HX-TIM3-5** | Z352412242 | TIM-3 | O=C(CC=1C=CC(F)=CC1)NC2=NNC=3C=CC=CC23 | 0.462 |
| **HX-TIM3-6** | Z1849736848 | TIM-3 | CCC(=O)NC1CCCN(C1)C(=O)[C@H]2C[C@]32CCOC4=CC=C(F)C=C43 | 0.884 |
| **HX-TIM3-7** | Z15905195 | TIM-3 | COC=1C=CC(=CC1OC)N2C(=NN=C2C=3C=CC=CC3)SCC(N)=O | 0.906 |
| **HX-TIM3-8** | Z4543490482 | TIM-3 | CC1=NC2=NC=NN2C(NCC=3C=C(NN3)C(=O)O)=C1C | 1.41 |
| **HX-TIM3-9** | Z444335508 | TIM-3 | CC1=CC=C(CCNC(=O)C2=CC=C(NC2=O)C=3C=CC=CC3)O1 | 1.66 |
| **HX-TIM3-10** | Z815667156 | TIM-3 | CC(C)CC=1C=CC(=CC1)S(=O)(=O)NC=2C=CC(=O)N(C)C2 | 1.71 |
| **HX-TIM3-11** | Z2222577366 | TIM-3 | CC=1N=CC=C(CCC(=O)N2CCCC(C2)C3=NOC(=N3)C4CC4)N1 | 1.75 |
| **HX-TIM3-12** | Z2007133546 | TIM-3 | CC1=CC=2C(=O)NC(CC=3C=CC(=CC3C)[N+]([O-])=O)=NC2S1 | 3.28 |
| **HX-TIM3-13** | Z1748892462 | TIM-3 | CC1CN(CCN1CCO)C(=O)CCOC=2C=CC(Cl)=CC2[N+]([O-])=O | 4.2 |
| **HX-TIM3-14** | Z57244559 | TIM-3 | CCOC(=O)C1=C(C)NC(NC(=O)NC=2C=C(C)C=CC2C)=C1C | 5.85 |
| **HX-TIM3-15** | Z92654463 | TIM-3 | FC=1C=CC(=CC1)C2=CN=C(CSC3=NN=NN3C4=CC=CC(F)=C4)O2 | 6.39 |
| **HX-TIM3-16** | Z2864024257 | TIM-3 | CN1C=CC(=N1)N2C=C(N=N2)C3=CSC(=N3)N4CCOCC4 | 7.12 |
| **HX-TIM3-17** | Z2482845318 | TIM-3 | CN(CC=1C=CC=2OCCOC2C1)C(C(=O)O)C=3C=CC=CC3F | 8.99 |
| **VISTA Confirmed Binders (n = 13)** | | | | |
| **HX-VISTA-1** | Z220421602 | VISTA | C1=NN(C=2C=CC=CC2)C3=NC=NC(NCC=4C=CC=CC4CN5CCOCC5)=C13 | 0.345 |
| **HX-VISTA-2** | Z1241130727 | VISTA | N#CC=1C=CC=CC1OCCN2CCN(CC3=CC(=O)N4C=CSC4=N3)CC2 | 0.931 |
| **HX-VISTA-3** | Z400997348 | VISTA | COC=1C=CC=CC1NC(=O)C2CCN(CC(=O)NCC=3C=CC(F)=CC3)CC2 | 1.01 |
| **HX-VISTA-4** | Z2451675322 | VISTA | COC=1C=CC(=CC1OC)CCN2CC(CC2=O)C(=O)NC3CCCNC3C | 1.24 |
| **HX-VISTA-5** | Z198290288 | VISTA | CC(C)(C)NC(=O)CNC(=O)C=1C=CC=2NN=NC2C1 | 1.37 |
| **HX-VISTA-6** | Z2045752765 | VISTA | OCCC1CCCCCN1CC2=CN3C=CC=NC3=N2 | 1.81 |
| **HX-VISTA-7** | Z838748444 | VISTA | CC1=NC(=CS1)COC=2C=CC=C(C2)C(=O)N(C)C3=CC=CC(C#N)=C3 | 1.83 |
| **HX-VISTA-8** | Z2199016863 | VISTA | CC1(C)CN(C(=O)NC=2C=CC=C(C2)C(N)=O)C1C3=CC=CN=C3 | 2.12 |
| **HX-VISTA-9** | Z1575682136 | VISTA | CC=1C=CC(=CC1)NC(=O)N2CCCCC3CCCCC32 | 2.23 |
| **HX-VISTA-10** | Z167586026 | VISTA | COC=1C=CC=C(C1)C(NC(=O)C=2C=NN(C)C2)C3=NC=CN3C | 2.78 |
| **HX-VISTA-11** | Z1426173894 | VISTA | O=C(C=1C=CN=CC1)N2CCC(C2)NCC=3C=CNC3 | 3.27 |
| **HX-VISTA-12** | Z226316098 | VISTA | O=C(COC=1C=CC=CC1)N(CC=2C=CC=CC2)CC3=NC=4C=CC=CC4C(=O)N3 | 9.05 |
| **HX-VISTA-13** | Z2087169547 | VISTA | CN1C=NC(=C1)C(=O)N2CCOCC2C3=NN=C(O3)C4=CC=CS4 | 12.83 |

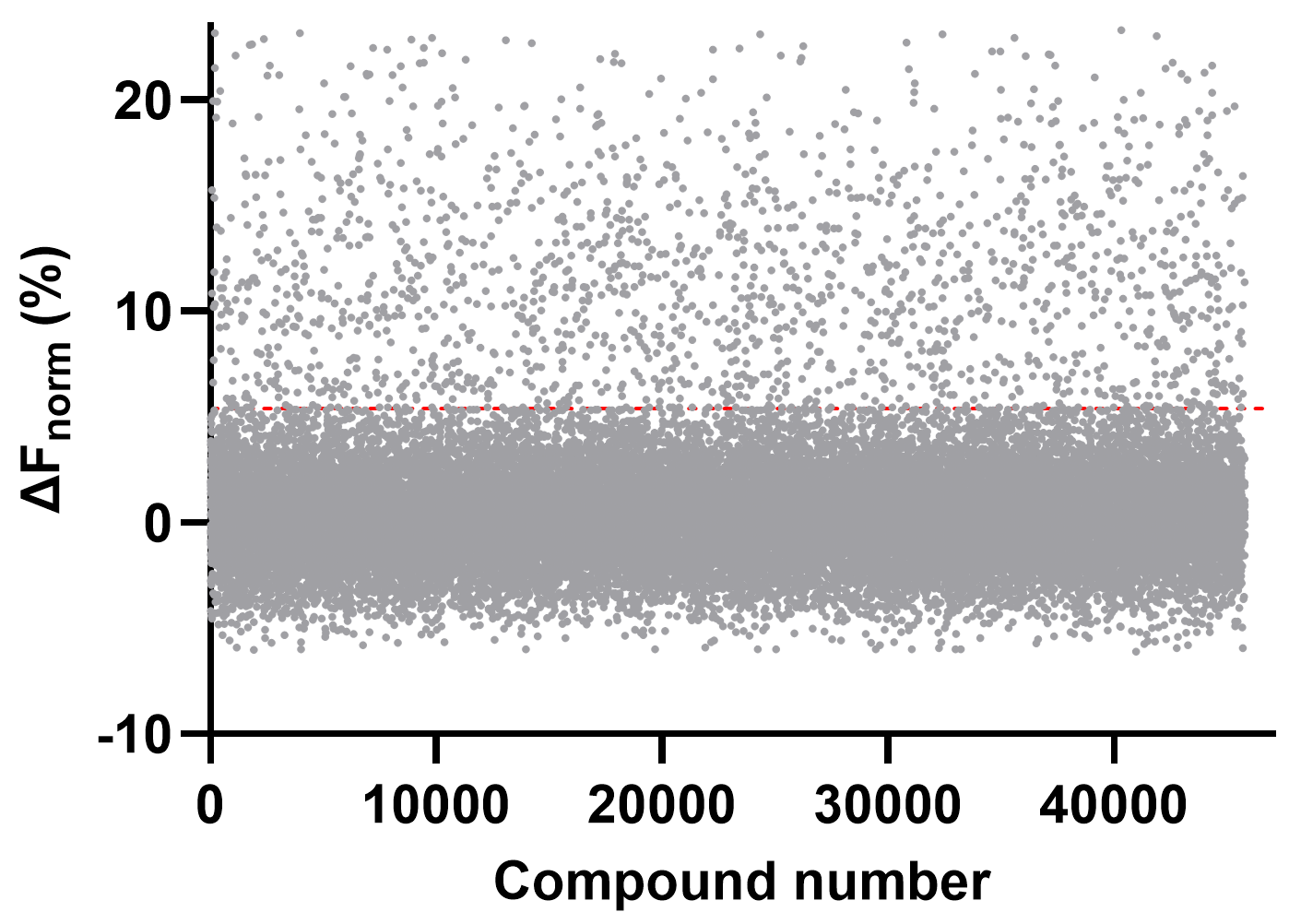

**Figure S1. Dianthus TRIC single-dose screening of the 45,760-compound Enamine training library against CD28.** Each dot represents one compound plotted by compound number vs. ΔF_norm_ (%). The red dashed line indicates the hit threshold.

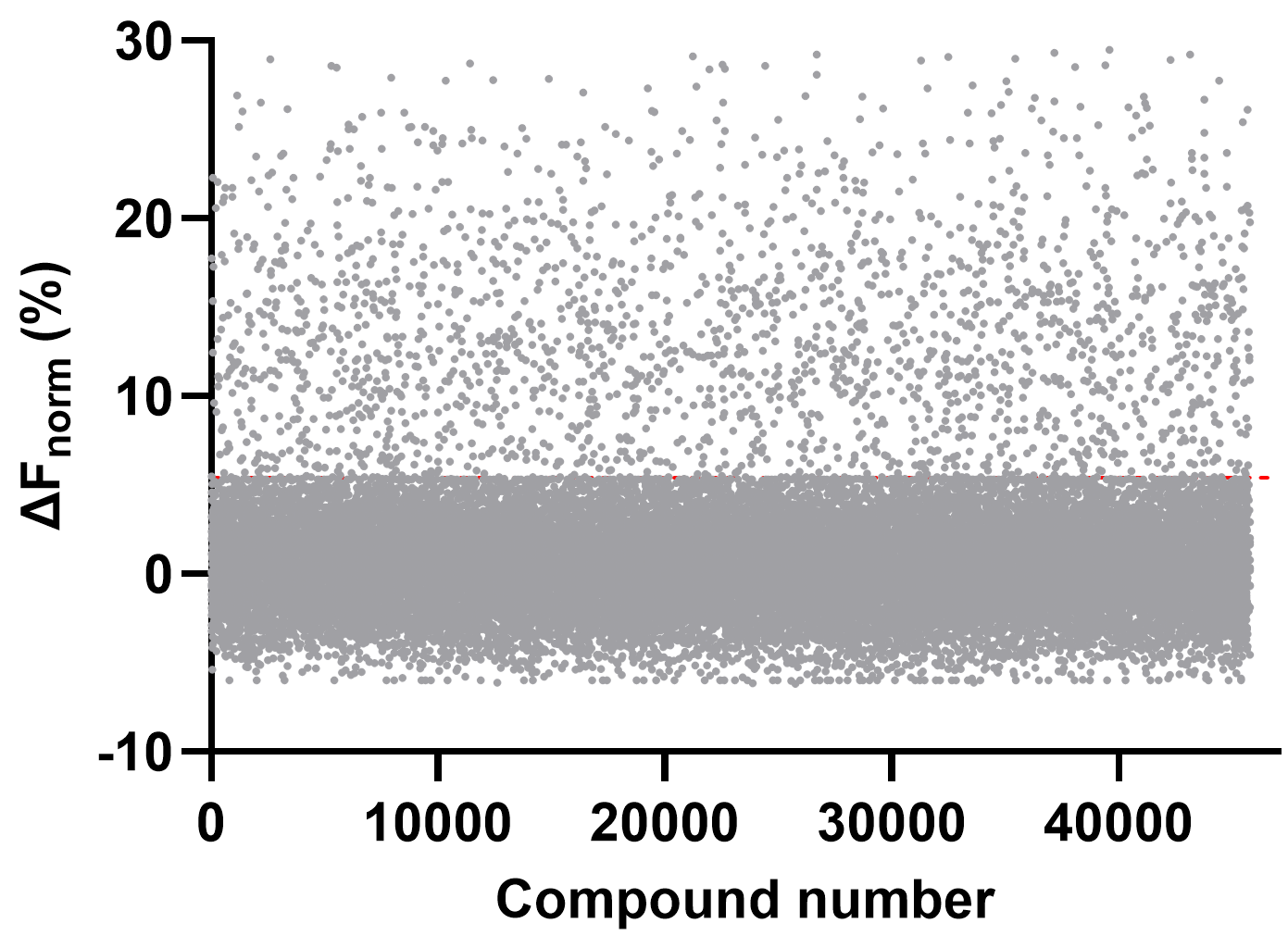

**Figure S2. Dianthus TRIC single-dose screening of the 45,760-compound Enamine training library against TIM-3.** Each dot represents one compound plotted by compound number vs. ΔF_norm_ (%). The red dashed line indicates the hit threshold.

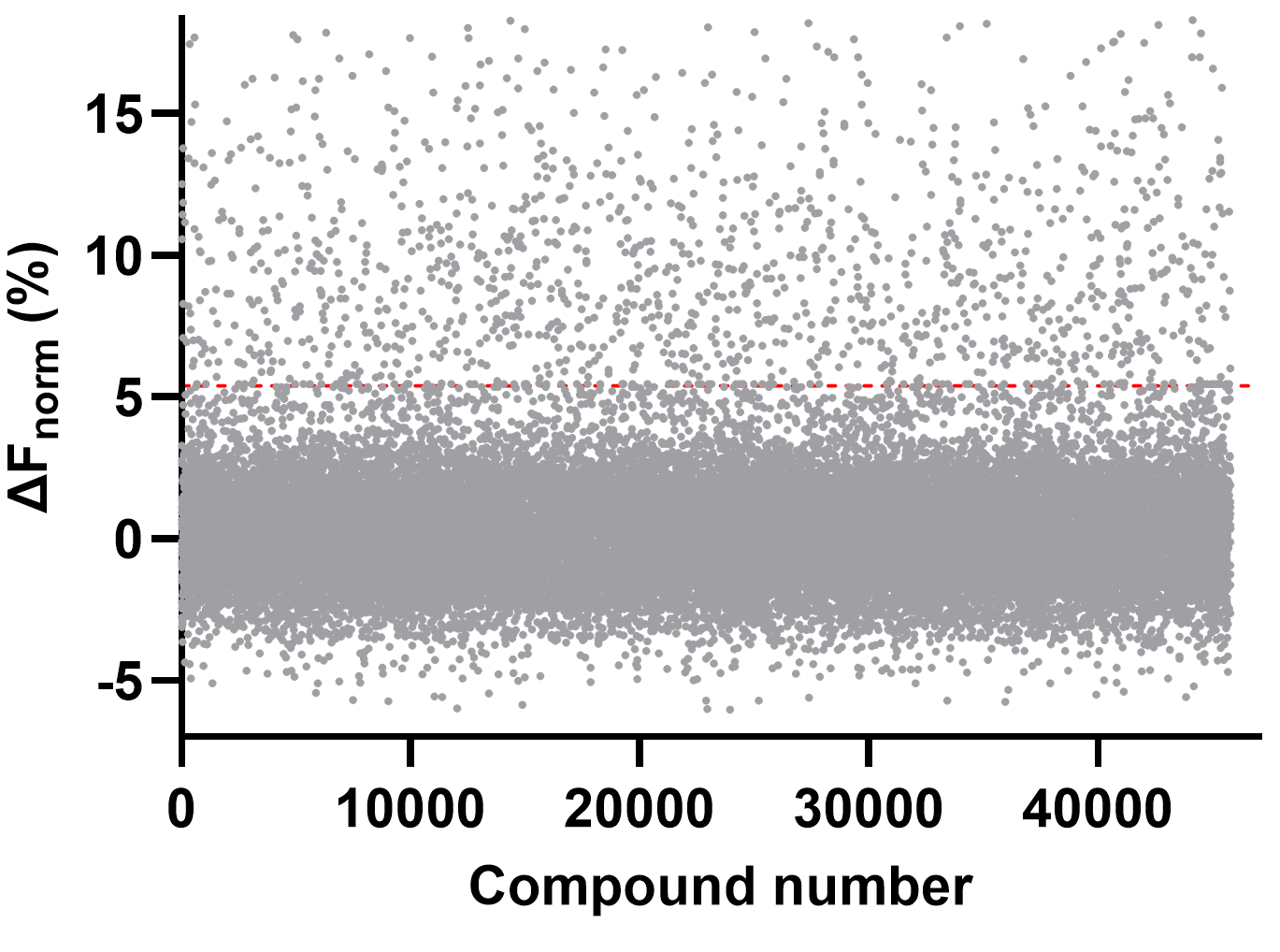

**Figure S3. Dianthus TRIC single-dose screening of the 45,760-compound Enamine training library against VISTA.** Each dot represents one compound plotted by compound number vs. ΔF_norm_ (%). The red dashed line indicates the hit threshold.

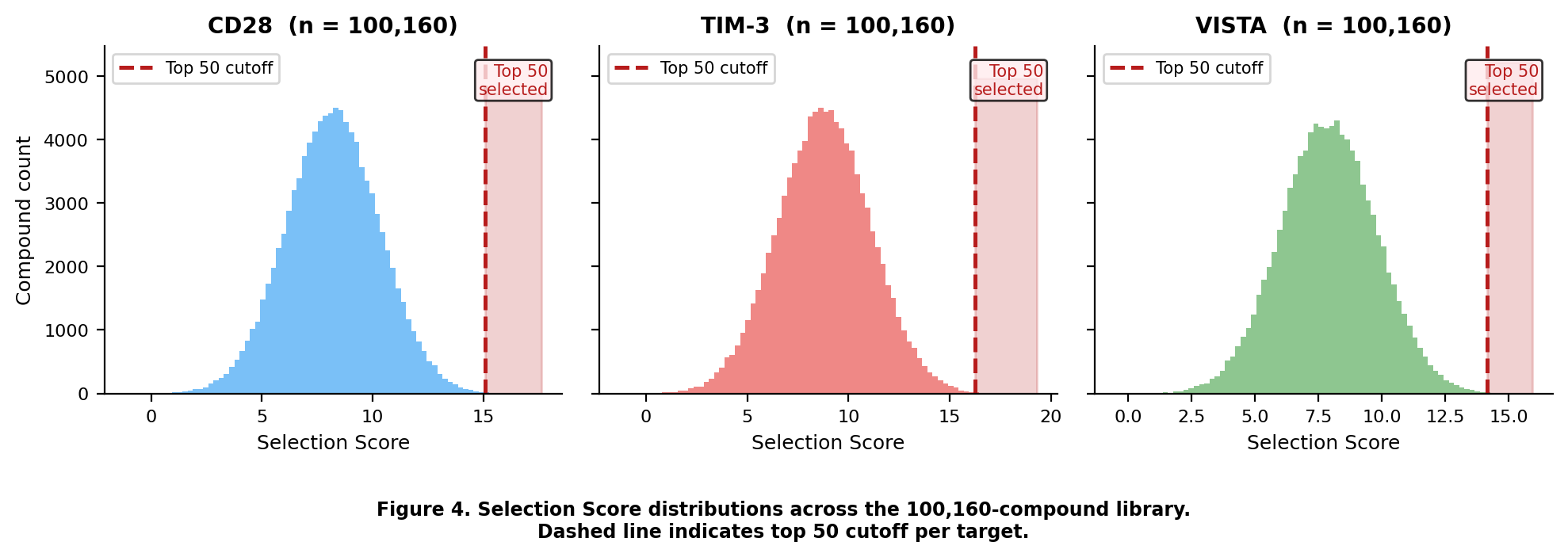

**Figure S4.** **Selection Score distributions across the 100,160-compound Enamine library for CD28, TIM-3, and VISTA.** Dashed red line and shaded region indicate the top 50 compound selection cutoff (99.95th percentile) per target. Scores reflect uncertainty-adjusted predicted ΔF_norm_ binding signals.
